# Comparisons between the Hydro Distillation and the Steam Distillation in the Extraction of Volatile Compounds from and the Anti-oxidative Activity of *Prunella Vulgaris*

**DOI:** 10.1101/2022.07.07.499219

**Authors:** William Chi Keung Mak

## Abstract

**Objective:** In this article, the aim is to verify a suggestion in our earlier study to explain the extraction dynamics of volatile compounds, being extracted from the herb *Prunella vulgaris* (PV) using the method of steam distillation. Then, the antioxidative property of PV is explored.

**Methods:** Because our earlier study suggested that the inefficient extraction using steam distillation was due to the mass of herb in the path of steam flow acting as an obstacle, we used hydro distillation which tried to eliminate this obstacle. We used gas chromatography – mass spectrometry (GC-MS) to characterize the volatile compounds extracted during the distillation process. Then, by treating the cancer cells from the cell line SCC154 with the distillate, the cancer cell cytotoxicity was assessed using the tetrazolium salt-based colorimetric test reagent, the Cell Counting Kit-8. The results provided the bases for comparisons. To assess the anti-oxidative activity of the PV distillate, Folin-Ciocalteu reagent was used.

**Results:** We successfully showed that the removal of the obstacle, formed by the mass of herb in the flow path of the uprising steam, enhanced the efficiency of volatile compound extraction and more volatile compounds could be extracted. Also, it was shown that the PV distillate did not exhibit anti-oxidative activity.

**Conclusions:** Hydro distillation is a more efficient method than steam distillation to extract volatile compounds from the PV herb. However, mild heating, which did not provide sufficient energy to the convection of the boiling water, did not move the floating herb on top of the boiling water; so, the obstacle still existed and limited the efficiency of extraction. For another issue of the antioxidant effect of the volatile compounds from PV, it was studied using the Folin-Ciocalteu reagent. It showed that the PV volatile compounds did not possess antioxidant property.

## 1. Introduction

*Prunella vulgaris* (PV) is a perennial low-growing plant which can be found in many regions in different continents worldwide^1^. In many cultures, Indians, Chinese, and Native Americans, preparations of the plant in the forms of salves, teas, decoction^2^ are used in the treatments of various minor ailments such as wounds and inflammation.

From a survey of literature, research works on PV extended back to 3 decades ago^3^. They can be categorised into three areas: the phytochemical area, the agricultural area, and the pharmacological area. In the phytochemical area, the attention was to investigate the extraction and analytical methods to isolate and identify the chemical constituents in PV^3,4,5,6^. In the agricultural area, PV is traditionally used in agriculture to feed domesticated livestock and used as veterinary medicine^7,8,9,10^. Investigations were also directed to explore the effects of the environmental conditions which affect the growth of PV. These included such factors as the water stress^11,12,13,14^, and soil fertilization^15,16,17^.

Most research efforts were, however, spent in the pharmacological area. Investigations were to explore the effects of PV on various pathogenic sources, such as the antimicrobial^18^, anti-viral^19,20,21,22,23,24,25^, anti-tumoral^26,27^, anti-inflammatory^28,29,30,31,32^, and others.

In the past, various bioactive compounds in PV were identified, including polysaccharides, polyphenolics and triterpenes^33^. Their biological activities were extensively studied in various research projects. Anti-tumorous, anti-inflammatory, antibiotic activities were found in polysaccharides^34^. Polyphenolics were found to have similar activities, while they also show anti-oxidative and anti-viral activities^35^. Among the triterpenes found in PV, betulinic acid and ursolic acid were shown to exhibit strong anti-allergic and anti-inflammatory effects when tests were conducted using cultured murine macrophages RAW 264.7 cells^36^. Rosmarinic acid, which is the compound used to identify PV^37^, was shown to have inhibitory effect of metastasis of breast and colon carcinoma^38,39^. PV extract and rosmarinic acid were shown to prevent UVB-induced damage and reduce oxidative stress by the reduction of ROS (reactive oxygen species)^40^.

Among research efforts in PV, very few efforts were spent on the investigation of volatile organic compounds (VOCs), however. The reason may be that the traditional method to prepare medicinal treatment for patients is by decoction (the boiling of herbs in water, which extracts the chemical ingredients into water). In this process, all volatile compounds evaporate away; thus, less attention has been given to the volatile compounds in previous studies. The few research efforts on PV VOCs concentrated on the analysis to identify the chemical composition in the plant^41,42,43^.

In this project, we used distillation to extract the volatile compounds from PV. In a previous work of this project^44^, we reported on the dynamics of how the volatile compounds were extracted during the whole distillation process. Using GS-MS as the analysis tool, we found that most volatile compounds came out continuously without much depletion even after the distillation progressed for a long time (over a half day which was the longest duration that the experiments could be carried out). Closer look showed that some rarer compounds did deplete early, however. They were only found in the portions at the very beginning (distillate was collected successively in 50 ml portions). Cell viability tests showed that the PV distillate had a weak cytotoxic activity against cancer cells SCC154, in a dosage dependent manner. This cytotoxicity also remained about the same for different portions, collected during the whole distillation process. This hinted that the cancer cell cytotoxicity was due to the less volatile compounds present in the distillate, since the most volatile compounds lost early during the initial stage of distillation.

Another article resulted from this project^45^ investigated the aging effect of the PV distillate. It showed that the aging of the distillate (for as long as 8 weeks) did not alter much the cell cytotoxicity of the distillate on the SCC154 cancer cells. This also again showed that the cell cytotoxicity should be due to the less volatile compounds which endured long after they were produced.

In the first part of this article, we explore the difference between the two distillation processes: steam distillation and hydro distillation. The purpose of doing this comparison was to verify a postulate we put forward in the previous article^44^ which explained the dynamics of the steam distillation process: The mass of herb hanging in the way of the passing steam from the boiling water below impeded the steam flow. Steam with the volatile compounds extracted re-condensed and dropped back to the boiling water below. This reduced the efficiency of the extraction of volatile compounds from the herb.

In the second part of this article, we report the exploration of the anti-oxidative effect of the PV herb.

## 2. Material and Methods

### 2.1 Materials

Dried spica of PV, imported from China, was obtained locally from the Traditional Chinese Medicine Clinic in the University of Technology Sydney in 2020. To be used as a reference standard in the authentication procedure, which was described previously^44^, rosmarinic acid purchased from Sigma Aldrich (R4033-10MG lot # BCBV7877) was used. The spica of the herb was finely pulverized using a grinder produced by Russell Hobbs (Classic chopper RHMFP2). GC-MS was carried out using a machine manufactured by Agilent Technologies (6390N Network GC System and 5973 Network Mass Selective Detector).

Tetrazolium-based colorimetric test reagent, which was used to assess the cell viability, Cell Counting Kit-8, was purchased from Sigma-Aldrich (product number: 96992-500-F). The cancer cell line which was used to assess the anti-tumoral effect was the SCC154 line. It was obtained internally in the university. To assess the anti-oxidative activity of the PV distillate, Folin-Ciocalteu (FC) reagent, purchased also from Sigma-Aldrich (product number: 47641-100-F), was used.

### 2.2 Methods

#### 2.2.1 The Distillation Processes

To assess the differences, two different distillation processes were compared. The steam distillation process was described previously^44^. Briefly, a two-ended reservoir flask containing the PV herb was placed above another flask, with water boiling in it. The herb was, thus, in the way of the rising steam flow. Steam carrying the volatile compounds extracted was condensed in a condenser on the other leg of the setup. The distillate was then collected during the whole process in 50-ml portions. The setup of the hydro distillation process was similar, except that the PV herb was submerged in the boiling water instead; so, the herb-containing flask was omitted. Steam from the boiling mixture was guided to the other leg of the setup and condensed by a condenser. Then, the distillate was collected in successive 50-ml portions during the distillation processes, in the same manner as in the steam distillation case.

For clarity, both setups for the steam distillation and for the hydro distillation are drawn side-by-side in Fig. 1 for comparison.

**Fig. 1.**
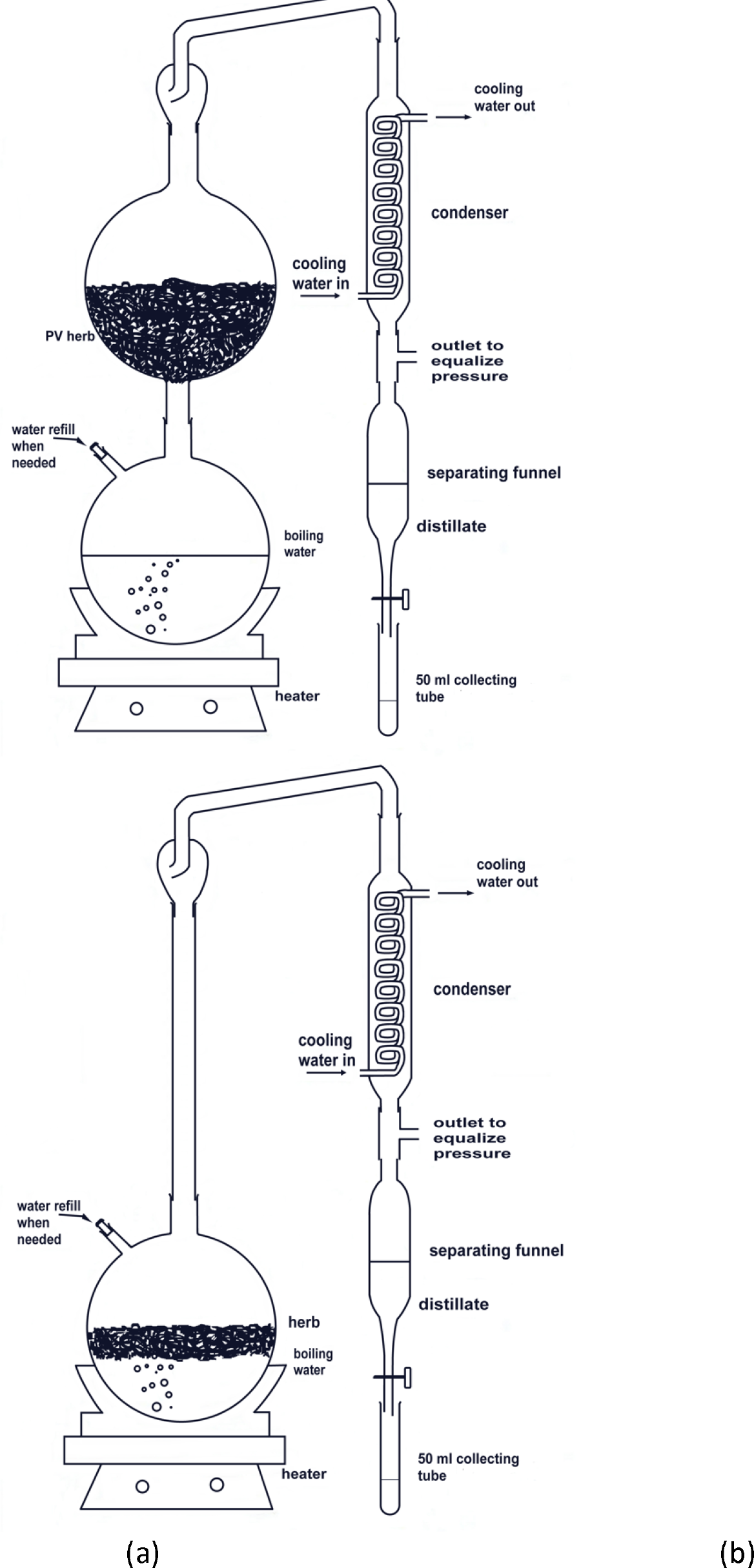
Both setups for (a) the steam distillation and (b) the hydro distillation processes are shown for comparison. The two-ended flask which contained the PV herb in the steam distillation setup was replaced by a long glass tube, to maintain the same potential energy which had to be overcome by the rising distillate-carrying steam. In the hydro distillation setup, the PV herb was submerged in the boiling water.

#### 2.2.2 GC-MS

The GC-MS machine is an Agilent machine, 6390N Network GC System and 5973 Network Mass Selective Detector. The capillary column in the gas chromatographer had dimensions of 30 m x 0.25 mm x 0.25 µm. The settings were as follows: Split mode injection; helium was used as the carrier gas at a flow rate of 1.2 mL/min under a pressure of 6.57 psi and an average velocity of 31 cm/s. At the inlet, the heater setting was at 250° C, at a flow rate of 27.7 mL/min and a pressure of 4.24 psi. The temperature profile of the oven is: Temperature held at 50° C for 3 min; then, ramping up to 250° C at a rate of 10° C/min; holding at 250° C for 3 min; then, ramping up to 280° C, where the temperature was at hold for a further 5 min, thus, making the total oven time of 34 minutes.

The mass spectrometer was set at electron impact mode, with a scan range between 30-500 amu, with a data rate of 20 Hz, and the detector set point at 280° C. The temperatures at the MS source and quad, were at 230 and 150° C respectively.

The chemical compounds in the distillate were partitioned from the aqueous medium into ethyl acetate before the GC-MS analysis was done to prevent the damage to the coating of the column inside the gas chromatographer.

Identifications of the various chemical constituents in the extracts were done by comparing the mass spectra obtained from the GC-MS runs of the samples, with those of known standard compounds obtained from the university chemistry laboratory. Comparisons were also made with the mass spectrum library of the National Institute of Standards and Technology (NIST08), available internally in the analysis software of the GC-MS machine.

#### 2.2.3 Cell Viability Test

The characterization of the cytotoxic effect of the PV distillate on the cancer cell lines SCC154 was done using the Cell Counting Kit-8, which probed the metabolic activities of the cancer cells using a tetrazolium salt WST-8, by a change of color of the salt, and thus indicated the cell viability. This is thus a colorimetric test to quantify the cell viability by measuring the absorbance at 450 nm.

We followed the procedures as stipulated in the document ‘Product Information’ provided by the vendor and was described earlier^44^. Experiments were repeated and data were taken in triplicate sets.

#### 2.2.4 Anti-oxidative test

The assessment of the anti-oxidative activity of the PV herb was done using the FC reagent, purchased from Sigma-Aldrich. The FC assay is a popular standardized method in the measurement of anti-oxidative capacity of food products and dietary supplements^46^. The procedure to conduct such a test was adopted from what was proposed in an article by Ainsworth & Gillespie^47^. Briefly, the procedure was as follows: 100 µL of the distillate was added to a 2-ml microtube. 200 µL FC reagent was added and then vortexed thoroughly. 800 µL 700 mM sodium carbonate was added and allowed to incubate at room temperature for 2 hours. 200 µL of the sample was transferred from the assay tube to a 96-well microplate. Three samples were repeated each, and thus, tests were done in triplicate sets. The absorbance of each well was measured by a microplate reader around 765 nm.

## 3. Results

The results can be divided into two parts: The first part looked at the differences in abundances of PV volatile compounds extracted during the two different distillation processes. This enables the study of the dynamics of the extraction processes and verifies the postulate in the previous article^44^ which explained the dynamics involved. The second part looked at the cell viability of the cancer cells SCC154 after they were treated by the PV distillate from the two different distillation processes.

### 3.1 Abundances of volatile compounds extracted

To compare between the two distillation methods, the settings of both methods should be adjusted to give meaningful comparison. This was done by adjusting the temperature of the heater so that the steam flow rates were the same. This was monitored by distillate collection rate to about 50 ml every 40 minutes. Four alkanes with different molecular weights were observed: decane, dodecane, tetradecane and hexadecane. They were chosen because their identities were positively identified^44^. They were also found to come out of the herb continuously with about constant rates even after the distillation process continued for a long time.

Fig. 2 shows the results.

**Fig. 2.**
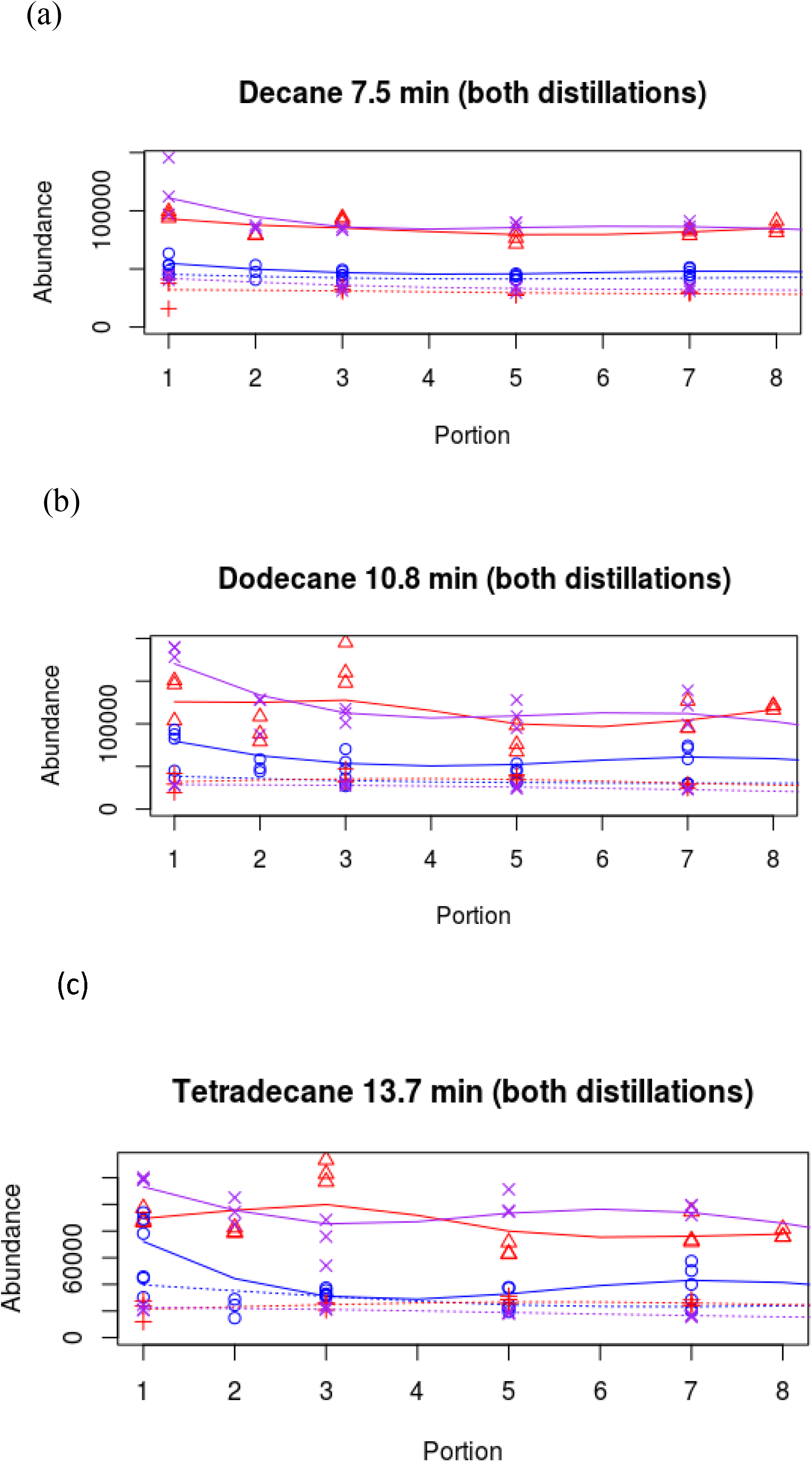
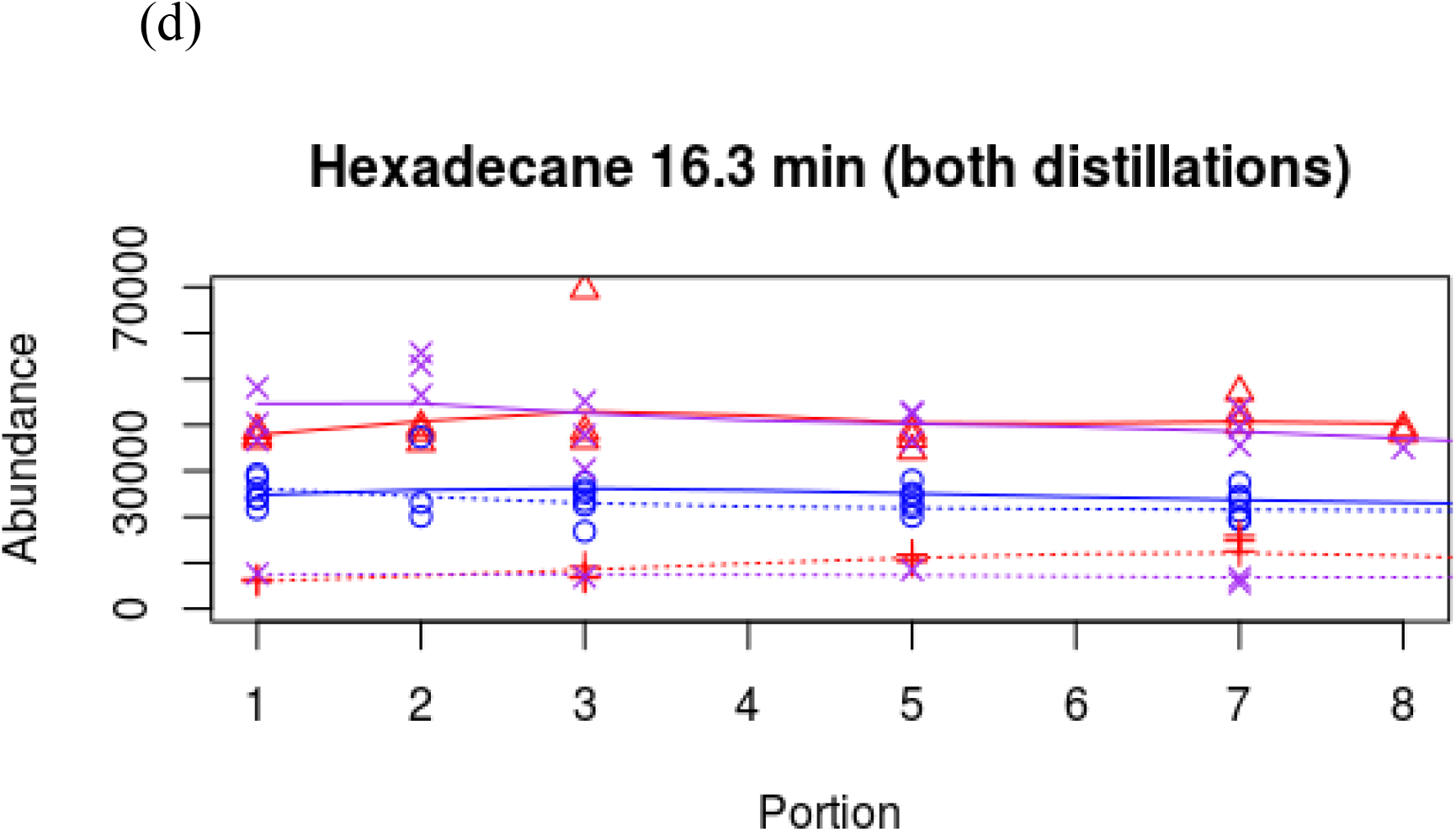
Abundances of alkanes extracted for the first 8 portions of 50 ml each. The dotted lines pertain to steam distillation, and the solid lines pertain to the hydro distillation. Three different amounts of PV herb were used: 15 g (blue), 25 g (red), 35 g (purple). The numbers appear in the sub-titles are the retention time of the compounds in minutes in the GC-MS coil. The smoothing lines were fitted to the data using cubic smoothing splines with a degree of smoothing parameter equal to 0.4.

With respect to steam distillation, the results repeated what was found in the previous article^44^. The amounts of volatile compounds extracted reached the optimal when 15 g of PV herb was used. It was postulated that, as more herb was used, the mass of herb in the way of steam flow became more and more obstructive and more volatile compounds were re-condensed and dropped back to the boiling liquid below. For the case of hydro distillation, the amounts of distillate obtained were larger through the whole distillation process than steam distillation at the same steam flow rate. This agrees well with the suggestion that the mass of PV herb hanging in the path of the steam flow acts as an obstacle in the case of steam distillation process. Moreover, the amount of 15 g of PV herb was no longer the optimal value to get the maximum amounts of volatile compounds, increasing the amount of herb to 25 g and 35 g extracted more volatile compounds. There was less resistance to the steam flow.

A closer and careful observation of Fig. 2 shows that, however, an increase of herb from 25 g to 35 g, however, did not increase the amounts of volatile compounds obtained. This seemed to contradict the intuitive suggestion that the hydro distillation process eliminated the obstacle of a mass of herb in the steam flow path, and thus, enhanced the extraction efficiency. However, looking more meticulously as the distillation proceeded revealed the reason. Because within the experimental setup, it was a nearly completely closed space except a tiny opening on the condenser side to allow a balance of pressure inside and outside. This balance opening was, however, not enough if the heating and the steam pressure was immense. The buildup of pressure might cause the boiling liquid to gush out or even trigger an explosion. To ensure safety, the heating needed to be limited. A consequence of this was that the convection of the boiling liquid had to be limited and was weak. When only 15 g of herb was used, the convection could still raise some boiling liquid above the herb floating at the surface and carried some herb in the convection current. When 25 g and 35 g of herb were used, the floating layer of herb at the surface was so thick that the weak convection current could not move it, and only bubbling steam could get through. Thus, the situation became similar to that found in steam distillation: the layer of herb at the liquid surface posed an obstacle to the steam flow, and so, saturation of extracted volatile compounds was again observed.

### 3.2 Cancer cell cytotoxicity

Cancer cell cytotoxicity was tested using the Cell Counting Kit-8 purchased from Sigma-Aldrich. The kit provides a means to check the cell metabolic activity through a colorimetric method using the tetrazolium salt WST-8. The procedure followed that provided by the vendor and was discovered previously^44^. The cancer cell line used was SCC154. Fig. 3 shows the results.

**Fig. 3.**
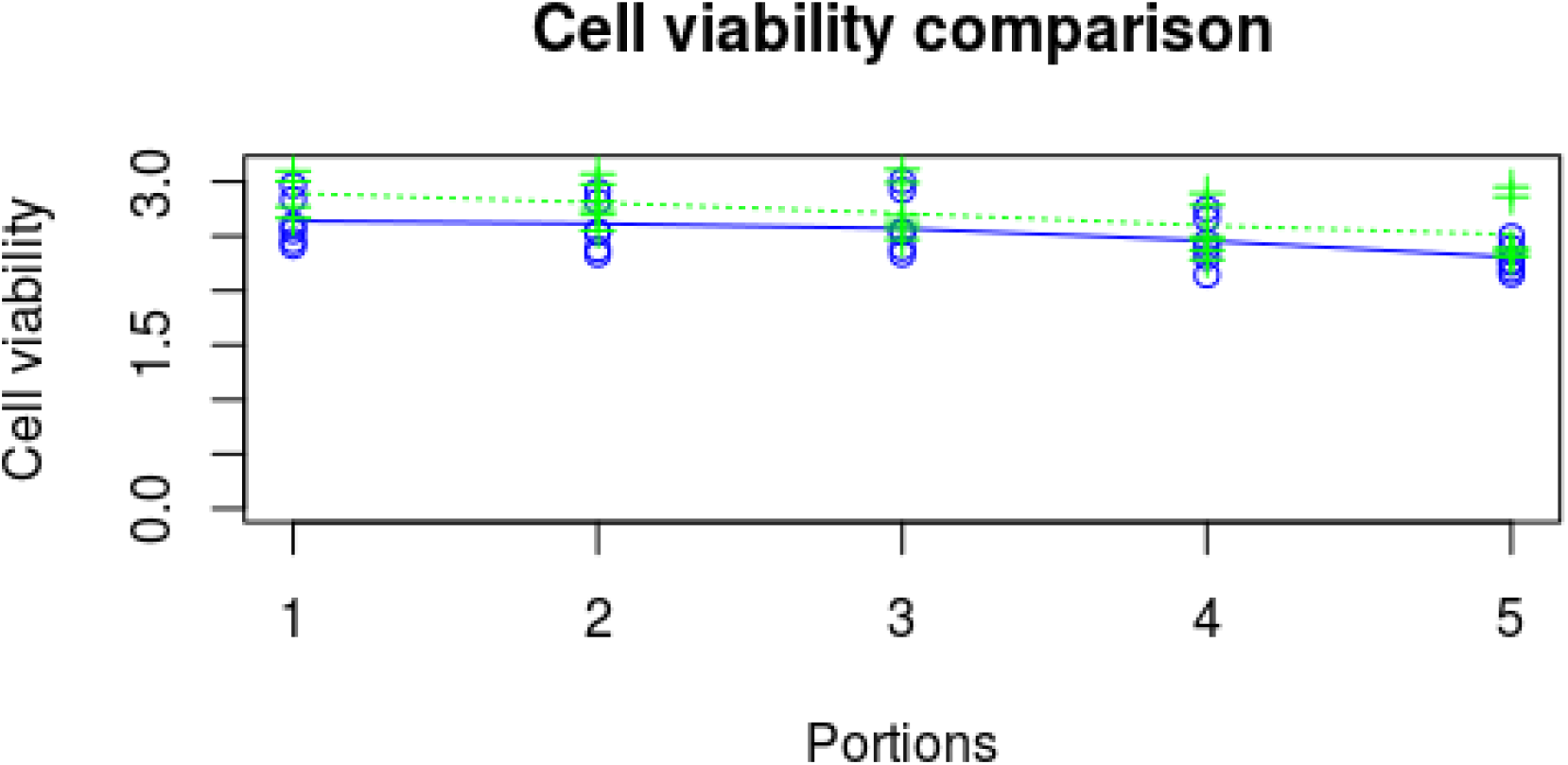
The cell viability of the cancer cells SCC154, treated by distillates from both hydro distillation (blue solid line and points) and steam distillation (green dotted line and points). The weights of herb used were 15 g. Only the first 5 portions collected during the distillation processes were used to plot the curves.

The cell viabilities found in this experiment agreed with what was found previously^44^. By visual observation of the curves, the cell cytotoxicity remained about the same for distillates in different portions obtained at different time during the distillation process, but was a little bit higher for later portions, which were collected later in the distillation processes. However, using the statistical two-way ANOVA method to analyze the differences among the portions gave a p-value of 0.0048. Using the commonly used threshold of p=0.05, the null hypothesis was rejected and thus there was significant differences in cell cytotoxicity among the portions.

One more observation is that the cell cytotoxicity of the distillate from the hydro distillation was higher than that obtained from the steam distillation. This result complied with what was found previously: Hydro distillation extracted more volatile compounds; the volatile compounds were more concentrated and thus the cell cytotoxicity was higher. Again, applying the two-way ANOVA analysis on the difference in cytotoxicity between hydro and steam distillation, it gave a p-value of 0.0061. Thus, the null hypothesis was also rejected, and there is a significant difference in anti-tumorous activities of the distillates obtained from hydro and steam distillation.

### 3.3 Anti-oxidative activity of the PV distillate

FC reagent was used to check the anti-oxidative capability of the PV herbal distillate, according to the procedures described earlier.

However, measurement of the absorbance of the treated PV distillate at 765 nm by a microplate reader showed that there was no peaking at this wavelength. The measurement curve remained flat in the vicinity of 765 nm (Fig. 4). Using one-way ANOVA analysis gave a p-value of 0.994. This means that the null hypothesis was accepted. The PV herbal distillate did not show anti-oxidative activity.

**Fig. 4.**
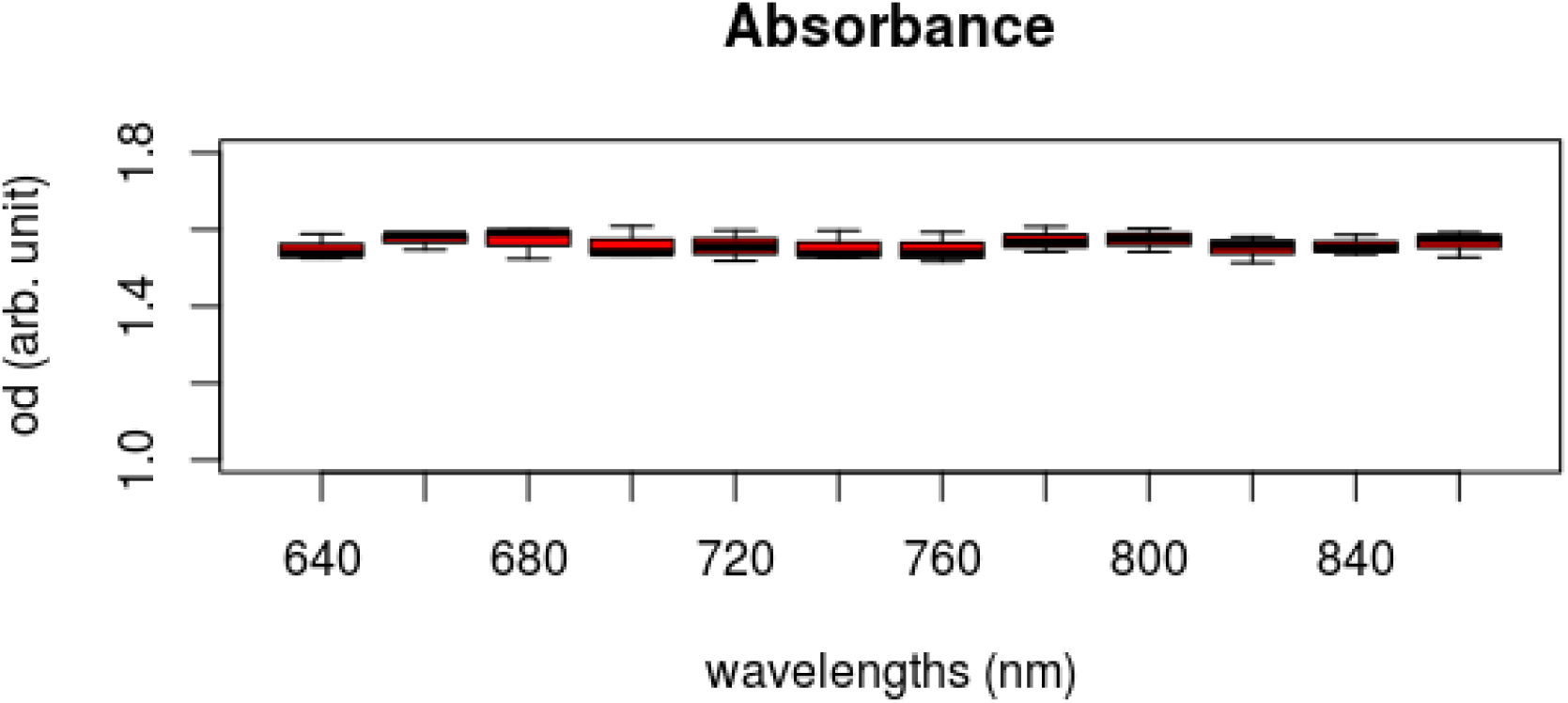
Absorbance measurements around 765 nm in Folin-Ciocalteu test

## 4. Conclusion

In this study, we compared the two distillation processes: the hydro distillation and the steam distillation. The main aim of this comparison is to verify a suggestion in a previous article^44^ which explained the dynamics of the steam distillation process. In that article, we found that the amounts of volatile compounds extracted did not increase with the amount of PV herb used when the amount of herb was more than 15 g. In fact, the amounts of volatile compounds obtained decreased as more herb was used. We postulated that as the mass of herb was put in the path of steam flow, it became an obstacle which re-condensed the volatile compounds, and they dropped back into the boiling water below. To test whether this postulate was valid, we removed this obstacle in the steam path by using hydro distillation. In this setup, the herb was submerged into the boiling water, and thus the obstacle in the steam path was removed. What we found experimentally validated this postulate. At the same steam flow rate, the amounts of volatile compounds extracted by hydro distillation were more than the corresponding amounts extracted by steam distillation. However, unexpectedly, the amounts of volatile compounds extracted still saturated and could not increase further when the amount of herb used was more than 25 g. Careful observation helped to discover that when the heating was mild to ensure safety, the convection of the boiling liquid was not strong enough to move the floating herb at the surface when it was thick, and this still became an obstacle in the steam path. Then, we compare the cancer cell cytotoxicity due to the two different distillation processes. We found that the distillate from hydro distillation was more cytotoxic to the cancer cells. This, once again, showed that the PV herb is cytotoxic to cancer cells, SCC154, in a dosage dependent manner. This is because we showed above that hydro distillation extracted more volatile compounds than steam distillation, and thus, the distillate was more concentrated.

The second part of this study looked at the anti-oxidative activity of PV herb, with the use of FC reagent. Disappointedly, no anti-oxidative activity was observed. This means that anti-oxidative compounds such as phenolics^28,47^ were not present in the distillate.

## 5. Discussion

The main aim in the study of this article is to demonstrate the dynamics of the steam distillation of PV herb is as postulated in the previous article^44^. The mass of herb hanging above the boiling water in the path of steam flow was an obstacle to hamper the efficiency of the extraction of volatile compounds. The hydro distillation is a process which eliminates this obstacle. This study demonstrates that the hydro distillation process achieved a higher efficiency than the steam distillation process; more volatile compounds were obtained at the same steam flow rate. On the other hand, looking at the different cancer cell cytotoxicity due to treatments using distillates from the two distillation processes revealed that the distillate from hydro distillate was more potent. This, once again, demonstrated the cancer cell cytotoxicity was dosage dependent because the distillate from hydro distillation was more concentrated in volatile compounds. An unexpected observation in this study, however, was that the volatile compounds extracted did not increase without limit when more and more herb was used for hydro distillation. Careful observation disclosed that the mass of floating herb on the surface of the boiling water again presented as an obstacle to the steam flow. The heating was kept mild to assure safety and thus the convection of the boiling liquid underneath was not strong enough to move the floating herb on top when the layer was thick.

The observations above have implications to the production process, when we apply the results found here to the manufacture of medicine which makes use of the PV volatile compounds. The herbal distillation is a more efficient method than steam distillation. This is convenient because the current practice of producing drugs, say, in granule form, is by boiling the herb submerged in water inside a big tank. So, hydro distillation becomes just a small additional step to collect the steam with volatile compounds above the boiling liquid. On the other hand, the use of steam distillation implies a different setup of equipment and thus additional cost for drug manufacture. An additional advantage is that hydro distillation extracts more volatile compounds at the same steam flow rate, and thus more efficient.

However, the observation that mild heating (and thus a weak convection liquid current) does not eliminate the obstacle of unmoved herb in the steam flow path, which limits the extraction efficiency. So, the design of the manufacture process should keep the heating of the decoction mixture high enough to achieve a better extraction efficiency, by ensuring the convection current carries the herb with it. However, in a previous paper^45^, it was shown that the composition of the extracted volatile compounds did change with the heating temperature. So, there needs further investigation in the process design to get the optimal settings. Previous studies^44,45^ also pointed to the fact that, as far as anti-tumoral activity is concerned, there are only very relaxed requirements on when the volatile compounds are taken, because volatile compounds taken at different time during the distillation process or taken many days afterward do not differ much in their potency. This offers much convenience to the design of the manufacture process.

The investigation into the anti-oxidative activity of PV volatile compounds showed that they did not consist of any anti-oxidative components. Many previous studies^4,31,48,49,50,51^ showed that non-volatile compounds in PV possess anti-oxidative property. So, just the distillation did not extract these anti-oxidative compounds as they are not volatile. Particularly, phenolics such as hydroxycinnamic acids, flavonoids, anthocyanins, tannins^47^ were not present in the distillate, and they were not responsible for the anti-tumorous property of the distillate.

One issue which may warrant further investigation is on the encapsulation of the VOCs extracted. Some research efforts were reported previously, such as the use of polymer-based microcapsules^52,53^. Practical applications were reported^54,55^ to treat herpes simplex viral disease using the VOCs from PV. Cream, which locked the VOCs from loss, was made by incorporating the essential oils from PV, and then applied topically on infected skin areas.

Finally, unlike Western pharmacology which identifies and singles out a particular phytochemical compound to be used as a drug, the practice of Chinese medicine is to make use of the synergistic effects of all constituent compounds in an herb or a combination of many herbs being used together. So, we just looked at the anti-tumoral activity of the volatile compounds in the distillate as a whole and intentionally avoid identifying individuals. So, identification of the individual anti-tumorous volatile PV compounds may be another direction of future works.

## List of Abbreviations

PV: *Prunella vulgaris*
VOC: volatile organic compound
GC-MS: gas chromatography – mass spectrometry
ROS: reactive oxygen species
NIST: National Institute of Standards and Technology
ANOVA: Analysis of variance

## Statements and Declarations

- There exist no competing interests which may inappropriately influence the author’s actions and the integrity of the research reported.
- There are no financial interests, which may be gained or lost from publication of the article. There is no financial gain, such as consulting fees or other remuneration from the publication of the article.
- The university which the author affiliated to has no financial gain or loss from the publication of the article.

## Acknowledgements

The first author wishes to thank his supervisors, Dr. Sean Walsh, and Prof. Alison Ung, for their guidance and support all along this research project.

